# Natural variation in teosinte at the domestication locus *teosinte branched1* (*tb1*)

**DOI:** 10.1101/005819

**Authors:** Laura Vann, Thomas Kono, Tanja Pyhäjärvi, Matthew B. Hufford, Jeffrey Ross-Ibarra

## Abstract

**Premise of the study:** The *teosinte branched1* (*tb1)* gene is a major QTL controlling branching differences between maize and its wild progenitor, teosinte. The insertion of a transposable element (*Hopscotch*) upstream of *tb1* is known to enhance the gene’s expression, causing reduced tillering in maize. Observations of the maize *tb1* allele in teosinte and estimates of an insertion age of the *Hopscotch* that predates domestication led us to investigate its prevalence and potential role in teosinte.

**Methods:** Prevalence of the *Hopscotch* element was assessed across an Americas-wide sample of 837 maize and teosinte individuals using a co-dominant PCR assay. Population genetic summaries were calculated for a subset of individuals from four teosinte populations in central Mexico. Phenotypic data were also collected using seed from a single teosinte population where *Hopscotch* was found segregating at high frequency.

**Key results:** Genotyping results indicate the *Hopscotch* element is found in a number of teosinte populations and linkage disequilibrium near *tb1* does not support recent introgression from maize. Population genetic signatures are consistent with selection on this locus revealing a potential ecological role for *Hopscotch* in teosinte, but a greenhouse experiment does not detect a strong association between *tb1* and tillering in teosinte.

**Conclusions:** Our findings suggest the role of *Hopscotch* differs between maize and teosinte. Future work should assess *tb1* expression levels in teosinte with and without the *Hopscotch* and more comprehensively phenotype teosinte to assess the ecological significance of the *Hopscotch* insertion and, more broadly, the *tb1* locus in teosinte.

## INTRODUCTION

Domesticated crops and their wild progenitors provide an excellent system in which to study adaptation and genomic changes associated with human-mediated selection (Ross-Ibarra et al., 2007). Plant domestication usually involves a suite of phenotypic changes such as loss of seed shattering and increased fruit or grain size, which are commonly referred to as the ‘domestication syndrome’ (Olsen and Wendel, 2013), and much of the study of domestication has focused on understanding the genetic variation underlying these traits (Olsen and Gross, 2010). Because most domesticates show reduced genetic diversity relative to their wild counterparts, effort has been made to identify agronomically useful variation in crop wild relatives (Flint-Garcia et al., 2009). Often, after identification, the alleles conferring these beneficial traits are bred into domesticates for crop improvement. For example, *Oryza rufipogon*, the wild progenitor of domesticated rice, has proven useful for the integration of a number of beneficial QTL controlling traits such as grain size and yield into domesticated rice (Kovach and McCouch, 2008). In addition to researching the role of wild alleles in domesticates, researchers have also investigated the role of variation in domesticated taxa in the evolution of feral and weedy populations (Ellstrand et al., 2010). But even though domesticated alleles are often found segregating in wild relatives (Gallavotti et al., 2004; Sigmon and Vollbrecht, 2010), we know almost nothing about the ecological role of this variation in natural populations. In this paper we present an ecological genetic analysis of the domestication locus *tb1*, and specifically the domesticated haplotype at *tb1*, in natural populations of the wild ancestor of domesticated maize.

Maize (*Zea mays* ssp. *mays*) was domesticated from the teosinte *Zea mays* ssp. *parviglumis* (hereafter, *parviglumis*) roughly 9,000 B.P. in southwest Mexico (Piperno et al., 2009; Matsuoka et al., 2002). Maize and the teosintes are an attractive system in which to study domestication due to the abundance of genetic tools developed for maize and well-characterized domestication loci (Hufford et al., 2012a; Doebley, 2004; Hufford et al., 2012b). Additionally, large, naturally-occurring populations of both *parviglumis* and the highland teosinte *Zea mays* ssp. *mexicana* (hereafter, *mexicana*) can be found throughout Mexico (Wilkes, 1977; Hufford et al., 2013), and genetic diversity of these taxa is estimated to be high (Ross-Ibarra et al., 2009).

Many morphological changes are associated with maize domestication, and understanding the genetic basis of these changes has been a focus of maize research for a number of years (Doebley, 2004). One of the most dramatic changes is found in plant architecture: domesticated maize is characterized by a central stalk with few tillers and lateral branches terminating in a female inflorescence, while teosinte is highly tillered and bears tassels (male inflorescences) at the end of its lateral branches. The *teosinte branched1* (*tb1)* gene, a repressor of organ growth, was identified as a major QTL involved in branching (Doebley et al., 1995) and tillering (Doebley and Stec, 1991) differences between maize and teosinte. A 4.9 kb retrotransposon (*Hopscotch*) insertion into the upstream control region of *tb1* in maize acts to enhance expression of *tb1*, thus repressing lateral organ growth (Doebley et al., 1997; Studer et al., 2011). Dating of the *Hopscotch* retrotransposon suggests that its insertion predates the domestication of maize, leading to the hypothesis that it was segregating as standing variation in populations of teosinte and increased to high frequency in maize due to selection during domestication (Studer et al., 2011). The effects of the *Hopscotch* insertion have been studied in maize (Studer et al., 2011), and analysis of teosinte alleles at *tb1* has identified functionally distinct allelic classes (Studer and Doebley, 2012), but little is known about the role of *tb1* or the *Hopscotch* insertion in natural populations of teosinte. Previous studies have confirmed the presence of the *Hopscotch* in samples of *parviglumis*, *mexicana*, and landrace maize; however little is known about the frequency with which the *Hopscotch* is segregating in natural populations.

In teosinte and other plants that grow at high population density, individuals detect competition from neighbors via the ratio of red to far-red light. An increase in far-red relative to red light accompanies shading and triggers the shade avoidance syndrome, a suite of physiological and morphological changes such as reduced tillering, increased plant height and early flowering (Kebrom and Brutnell, 2007). The *tb1* locus appears to play an important role in the shade avoidance pathway in *Zea mays* (Lukens and Doebley, 1999) and other grasses (Kebrom and Brutnell, 2007) via changes in expression levels in response to shading. Lukens and Doebley (1999) introgressed the teosinte *tb1* allele into a maize inbred background and noted that under low density conditions plants were highly tillered but that under high density, plants showed significantly reduced tillers and grew taller. Based on these results we hypothesize that the *Hopscotch* (*i.e.*, the domesticated allele) at *tb1* may play a role in the ecology of teosinte, especially in high-density populations. In this study we aim to characterize the distribution of the *Hopscotch* insertion in *parviglumis*, *mexicana*, and landrace maize, and to examine the phenotypic effects of the insertion in *parviglumis*. We use a combination of PCR genotyping for the *Hopscotch* element in our full panel and sequencing of two small regions upstream of *tb1* combined with a larger SNP dataset in a subset of teosinte populations to explore patterns of genetic variation at this locus. Finally, we test for an association between the *Hopscotch* element and tillering phenotypes in samples from a natural population of *parviglumis*.

## MATERIALS AND METHODS

### Sampling and genotyping

We sampled 1,110 individuals from 350 accessions (247 maize landraces, 17 *mexicana* populations, and 86 *parviglumis* populations; ranging from 1-38 individuals per population, with an average of 11 individuals per population for *parviglumis* and *mexicana* and 2 individuals per landrace accession) and assessed the presence or absence of the *Hopscotch* insertion (Appendix 1 and Appendix 2, See Supplemental Materials with the online version of this article). DNA was extracted from leaf tissue using a modified CTAB approach (Doyle and Doyle, 1990; Maloof et al., 1984). We designed primers using PRIMER3 (Rozen and Skaletsky, 2000) implemented in Geneious (Kearse et al., 2012) to amplify the entire *Hopscotch* element, as well as an internal primer allowing us to simultaneously check for possible PCR bias between presence and absence of the *Hopscotch* insertion due to its large size (*∼*5 kb). Two PCRs were performed for each individual, one with primers flanking the *Hopscotch* (HopF/HopR) and one with a flanking primer and an internal primer (HopF/HopIntR). Primer sequences are HopF, 5’-TCGTTGATGCTTTGATGGATGG-3’; HopR, 5’-AACAGTATGATTTCATGGGACCG-3’; and HopIntR, 5’-CCTCCACCTCTCATGAGATCC-3’ (Appendix 3 and Appendix 4, See Supplemental Materials with the online version of this article). Homozygotes show a single band for absence of the element (*∼*300 bp) and two bands for presence of the element (*∼*5 kb, amplification of the entire element, and *∼*1.1 kb, amplification of part of the element), whereas heterozygotes show all three bands (Appendix 2, See Supplemental Materials with the online version of this article). Since we developed a PCR protocol for each allele, if only one PCR resolved well, we scored one allele for that individual rather than infer the diploid genotype. We used Phusion High Fidelity Enzyme (Thermo Fisher Scientific Inc., Waltham, Massachusetts, USA) and the following conditions for amplifications: 98*°*C for 3 min, 30 cycles of 98*°*C for 15 s, 65*°*C for 30 s, and 72*°*C for 3 min 30 s, with a final extension of 72*°*C for 10 min. PCR products were visualized on a 1% agarose gel and scored for presence/absence of the *Hopscotch* based on band size.

### Genotyping analysis

To calculate differentiation between populations (F_ST_) and subspecies (F_CT_) we used HierFstat (Goudet, 2005). These analyses only included populations (*n* = 32) in which eight or more chromosomes were sampled. To test the hypothesis that the *Hopscotch* insertion may be adaptive under certain environmental conditions, we looked for significant associations between *Hopscotch* frequency and environmental variables using the software BayEnv (Coop et al., 2010). BayEnv creates a covariance matrix of relatedness between populations and then tests a null model that allele frequencies in populations are determined by the covariance matrix of relatedness alone against the alternative model that allele frequencies are determined by a combination of the covariance matrix and an environmental variable, producing a posterior probability (*i.e.,* Bayes Factor; Coop et al. 2010). We used teosinte (ssp. *parviglumis* and ssp. *mexicana*) genotyping and covariance data from Pyhäjärvi et al. (2013) for BayEnv, with the *Hopscotch* insertion coded as an additional biallelic marker. SNP data from Pyhäjärvi et al. (2013) were obtained using the MaizeSNP50 BeadChip and Infinium HD Assay (Illumina, San Diego, CA, USA). Environmental data were obtained from www.worldclim.org and soil data were downloaded from the Harmonized World Soil Database (FAO/IIASA/ISRIC/ISSCAS/JRC, 2012) at www.harvestchoice.org. Environmental data represent average values for the last several decades (climatic data) or are likely stable over time (soil data) and therefore represent conditions important for local adaptation of our samples. Information from these data sets was summarized by principle component analysis following Pyhäjärvi et al. (2013).

### Sequencing

In addition to genotyping, we chose a subset of *parviglumis* individuals for sequencing. We chose twelve individuals from each of four populations from Jalisco state, Mexico (San Lorenzo, La Mesa, Ejutla A, and Ejutla B). For amplification and sequencing, we selected two regions approximately 600 bp in size from within the 5’ UTR of *tb1* (Region 1) and from 1,235 bp upstream of the start of the *Hopscotch* (66,169 bp upstream from the start of the *tb1* ORF; Region 2). We designed the following primers using PRIMER3 (Rozen and Skaletsky, 2000): for the 5’ UTR, 5-’GGATAATGTGCACCAGGTGT-3’ and 5’-GCGTGCTAGAGACACYTGTTGCT-3’; for the 66 kb upstream region, 5’-TGTCCTCGCCGCAACTC-3’ and 5’-TGTACGCCCGCCCCTCATCA-3’ (Appendix 1, See Supplemental Materials with the online version of this article). We used Taq polymerase (New England Biolabs Inc., Ipswich, Massachusetts, USA) and the following thermal cycler conditions to amplify fragments: 94*°*C for 3 min, 30 cycles of 92*°*C for 40 s, annealing for 1 min, 72*°*C for 40 s, and a final 10 min extension at 72*°*C. Annealing temperatures for Region 1 and Region 2 were 59.7*°*C and 58.8*°*C, respectively. To clean excess primer and dNTPs we added two units of Exonuclease1 and 2.5 units of Antarctic Phosphatase to 8.0 µL of amplification product. This mix was placed on a thermal cycler with the following program: 37*°*C for 30 min, 80*°*C for 15 min, and a final cool-down step to 4*°*C.

We cloned cleaned fragments into a TOPO-TA vector (Life Technologies, Grand Island, New York, USA) using OneShot TOP10 chemically competent *E. coli* cells, with an extended ligation time of 30 min for a complex target fragment. We plated cells on LB agar plates containing kanamycin, and screened colonies using vector primers M13 Forward and M13 Reverse under the following conditions: 96*°*C for 5 min; then 35 cycles at 96*°*C for 30 s, 53*°*C for 30 s, 72*°*C for 2 min; and a final extension at 72*°*C for 4 min. We visualized amplification products for incorporation of our insert on a 1% agarose TAE gel.

Amplification products with successful incorporation of our insert were cleaned using Exonuclease 1 and Antarctic Phosphatase following the procedures detailed above, and sequenced with vector primers M13 Forward and M13 Reverse using Sanger sequencing at the College of Agriculture and Environmental Sciences (CAES) sequencing center at UC Davis. We aligned and trimmed primer sequences from resulting sequences using the software Geneious (Kearse et al., 2012). Following alignment, we verified singleton SNPs by sequencing an additional one to four colonies from each clone. If the singleton was not present in these additional sequences it was considered an amplification or cloning error, and we replaced the base with the base of the additional sequences. If the singleton appeared in at least one of the additional sequences we considered it a real variant and kept it for further analyses.

### Sequence analysis

For population genetic analyses of sequenced Region 1 and sequenced Region 2 we used the Libsequence package (Thornton, 2003) to calculate pairwise F_ST_ between populations and to calculate standard diversity statistics (number of haplotypes, haplotype diversity, Watterson’s estimator 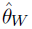, pairwise nucleotide diversity 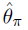, and Tajima’s D). To produce a visual representation of differentiation between sequences and examine patterns in sequence clustering by *Hopscotch* genotype we used Phylip (http://evolution.genetics.washington.edu/phylip.html) to create neighbor-joining trees with bootstrap-supported nodes (100 repetitions). For creation of trees we also included homologous sequence data from Maize HapMapV2 (Chia et al., 2012) for teosinte inbred lines (TILs), some of which are known to be homozygous for the *Hopscotch* insertion (TIL03, TIL17, TIL09), as well as 59 lines of domesticated maize.

### Introgression analysis

In order to assess patterns of linkage disequilibrium (LD) around the *Hopscotch* element in the context of chromosomal patterns of LD we used Tassel (Bradbury et al., 2007) and calculated LD between SNPs across chromosome 1 using previously published data from twelve plants each of the Ejutla A (EjuA), Ejutla B (EjuB), San Lorenzo (SLO), and La Mesa (MSA) populations (Pyhäjärvi et al., 2013). We chose these populations because we had both genotyping data for the *Hopscotch* as well as chromosome-wide SNP data for chromosome 1. For each population we filtered the initial set of 5,897 SNPs on chromosome 1 to accept only SNPs with a minor allele frequency of at least 0.1, resulting in 1,671, 3,023, 3,122, and 2,167 SNPs for SLO, EjuB, EjuA, and MSA, respectively. We then used Tassel (Bradbury et al., 2007) to calculate linkage disequilibrium (*r*^2^) across chromosome 1 for each population.

We examined evidence of introgression on chromosome 1 in these same four populations (EjuA, EjuB, MSA, SLO) using STRUCTURE (Falush et al., 2003) and phased data from Pyhäjärvi et al. (2013), combined with the corresponding SNP data from a diverse panel of 282 maize lines (Cook et al., 2012). SNPs were anchored in a modified version of the IBM genetic map (Gerke et al., 2013). Since STRUCTURE does not account for LD due to physical linkage we created haplotype blocks using a custom Perl script from Hufford et al. (2013, code available at http://dx.doi.org/10.6084/m9.figshare.1165577). In maize, LD decays over an average distance of 5500 bp (Chia et al., 2012); because LD decay is even more rapid in teosinte (Chia et al., 2012) we used a conservative haplotype block size of 5 kb. We ran STRUCTURE at K = 2 under the linkage model, performing 3 replicates with an MCMC burn-in of 10,000 steps and 50,000 steps post burn-in.

### Phenotyping of *parviglumis*

To investigate the phenotypic effects of the *Hopscotch* insertion in teosinte we conducted a phenotyping trial in which we germinated 250 seeds of *parviglumis* collected in Jalisco state, Mexico (population San Lorenzo; Hufford 2010) where the *Hopscotch* insertion is segregating at highest frequency (0.44) in our initial genotyping sample set. In order to maximize the likelihood of finding the *Hopscotch* in our association population we selected seeds from sites within the population where genotyped individuals were homozygous or heterozygous for the insertion. We chose between 10-13 seeds from each of 23 sampling sites. We treated seeds with Captan fungicide (Southern Agricultural Insecticides Inc., Palmetto, Florida, USA) and germinated them in petri dishes with filter paper. Following germination, 206 successful germinations were planted into one-gallon pots with potting soil and randomly spaced one foot apart on greenhouse benches. Plants were watered three times a day with an automatic drip containing 10-20-10 fertilizer, which was supplemented with hand watering on extremely hot and dry days.

Starting on day 15, we measured tillering index as the ratio of the sum of tiller lengths to the height of the plant (Briggs et al., 2007). Following initial measurements, we phenotyped plants for tillering index every 5 days through day 40, and then on day 50 and day 60. On day 65 we measured culm diameter between the third and fourth nodes of each plant. Following phenotyping we extracted DNA from all plants using a modified SDS extraction protocol. We genotyped individuals for the *Hopscotch* insertion following the PCR protocols listed above.

Tillering index data for each genotypic class did not meet the criteria for a repeated measures ANOVA, so we transformed the data with a Box-Cox transformation (*λ* = 0) in the Car Package for R (Fox and Weisberg, 2011) to improve the normality and homogeneity of variance among genotype classes. We analyzed relationships between genotype and tillering index and tiller number using a repeated measures ANOVA through a general linear model function implemented in SAS v.9.3 (SAS Institute Inc., Cary, NC, USA). Additionally, in order to compare any association between *Hopscotch* genotype and tillering and associations at other presumably unrelated traits, we performed an ANOVA between culm diameter and genotype using the same general linear model in SAS. Culm diameter is not believed to be correlated with tillering index or variation at *tb1* and is used as our independent trait for phenotyping analyses. SAS code used for analysis is available at http://dx.doi.org/10.6084/m9.figshare.1166630.

## RESULTS

### Genotyping

The genotype at the *Hopscotch* insertion was confirmed with two PCRs for 837 individuals of the 1,100 screened. Among the 247 maize landrace accessions genotyped, all but eight were homozygous for the presence of the insertion (Appendix 1 and Appendix 2, See Supplemental Materials with the online version of this article). Within our *parviglumis* and *mexicana* samples we found the *Hopscotch* insertion segregating in 37 (*n* = 86) and four (*n* = 17) populations, respectively, and at highest frequency within populations in the states of Jalisco, Colima, and Michoaćan in central-western Mexico (Fig. 1). Using our *Hopscotch* genotyping, we calculated differentiation between populations (F_ST_) and subspecies (F_CT_) for populations in which we sampled eight or more individuals. We found that F_CT_ = 0, and levels of F_ST_ among populations within each subspecies (0.22) and among all populations (0.23) are similar to genome-wide estimates from previous studies (Pyhäjärvi et al. 2013; Table 1). Although we found large variation in *Hopscotch* allele frequency among our populations, BayEnv analysis did not indicate a correlation between the *Hopscotch* insertion and environmental variables (all Bayes Factors *<* 1).

**Figure 1.**
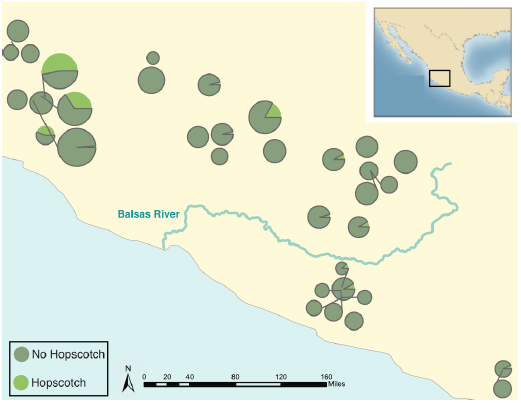
Map showing the frequency of the *Hopscotch* allele in populations of *parviglumis* where we sampled more than 6 individuals. Size of circles reflects number of alleles sampled. The Balsas River is shown, as the Balsas River Basin is believed to be the center of domestication of maize.

**Table 1.** Pairwise F_ST_ values from sequence and *Hopscotch* genotyping data

| **Comparison** | **Region 1** | **Region 2** | ***Hopscotch*** |
| --- | --- | --- | --- |
| EjuA & EjuB | 0 | 0 | 0 |
| EjuA & MSA | 0.326 | 0.328 | 0.186 |
| EjuA & SLO | 0.416 | 0.258 | 0.280 |
| EjuB & MSA | 0.397 | 0.365 | 0.188 |
| EjuB & SLO | 0.512 | 0.290 | 0.280 |
| MSA & SLO | 0.007 | 0 | 0.016 |

### Sequencing

To investigate patterns of sequence diversity and linkage disequilibrium (LD) in the *tb1* region, we sequenced two small (*<*1 kb) regions upstream of the *tb1* ORF in four populations. After alignment and singleton checking we recovered 48 and 40 segregating sites for the 5’ UTR region (Region 1) and the 66 kb upstream region (Region 2), respectively. For Region 1, Ejutla A has the highest values of haplotype diversity and *θ_π_*, while Ejutla B and La Mesa have comparable values of these summary statistics, and San Lorenzo has much lower values. Additionally, Tajima’s D is strongly negative in the two Ejutla populations and La Mesa, but is closer to zero in San Lorenzo (Table 2, Appendix 2, See Supplemental Materials with the online version of this article). For Region 2, haplotype diversity and *θ_π_*, are similar for Ejutla A and Ejutla B, while La Mesa and San Lorenzo have slightly lower values for these statistics (Table 2). Tajima’s D is positive in all populations except La Mesa, indicating an excess of low frequency variants in this population (Table 2). Pairwise values of F_ST_ within population pairs Ejutla A/Ejutla B and San Lorenzo/La Mesa are close to zero for both sequenced regions as well as for the *Hopscotch*, while they are high for other population pairs (Table 1). Neighbor joining trees of our sequence data and data from the teosinte inbred lines (TILs; data from Maize HapMapV2, Chia et al. 2012) do not reveal any clear clustering pattern with respect to population or *Hopscotch* genotype (Appendix 5, See Supplemental Materials with the online version of this article); individuals within our sample that have the *Hopscotch* insertion do not group with the teosinte inbred lines or domesticated maize that have the *Hopscotch* insertion.

**Table 2.** Population genetic statistics from resequenced regions near the *tb1* locus

| Population | # Haplotypes | Hap. Diversity | $\hat{\theta}_\pi$ | Tajima's D |
| --- | --- | --- | --- | --- |
| *Region 1 (5' UTR)* |  |  |  |  |
| EJUA | 8 | 0.859 | 0.005 | -1.650 |
| EJUB | 5 | 0.709 | 0.004 | -1.831 |
| MSA | 6 | 0.682 | 0.004 | -1.755 |
| SLO | 3 | 0.318 | 0.001 | -0.729 |
| *Region 2 (66kb upstream)* |  |  |  |  |
| EJUA | 8 | 0.894 | 0.018 | 0.623 |
| EJUB | 8 | 0.894 | 0.016 | 0.295 |
| MSA | 3 | 0.682 | 0.011 | -0.222 |
| SLO | 4 | 0.742 | 0.014 | 0.932 |

### Evidence of introgression

The highest frequency of the *Hopscotch* insertion in teosinte was found in *parviglumis* sympatric with cultivated maize. Our initial hypothesis was that the high frequency of the *Hopscotch* element in these populations could be attributed to introgression from maize into teosinte. To investigate this possibility we examined overall patterns of linkage disequilibrium across chromosome one and specifically in the *tb1* region. If the *Hopscotch* is found in these populations due to recent introgression we would expect to find large blocks of linked markers near this element. We find no evidence of elevated linkage disequilibrium between the *Hopscotch* and SNPs surrounding the *tb1* region in our resequenced populations (Figure 2), and *r*^2^ in the *tb1* region does not differ significantly between populations with (average *r*^2^ of 0.085) and without (average *r*^2^ = 0.082) the *Hopscotch* insertion. In fact, average *r*^2^ is lower in the *tb1* region (*r*^2^ = 0.056) than across the rest of chromosome 1 (*r*^2^ = 0.083; Table 3).

**Figure 2.**
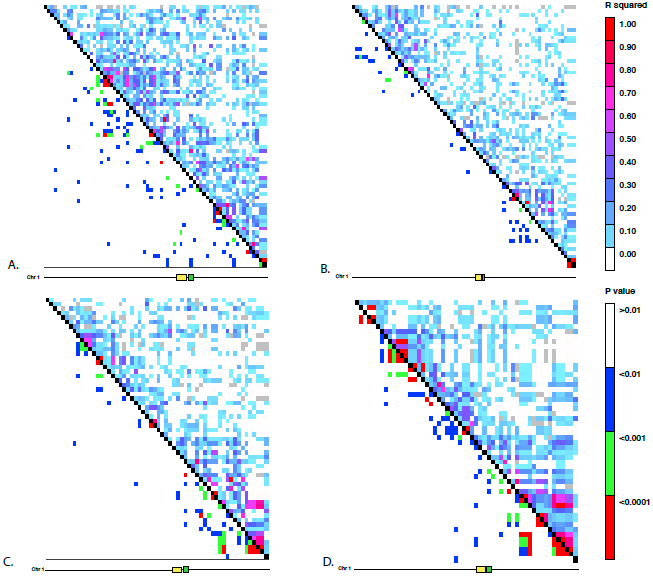
Linkage disequilibrium for SNPs in Mb 261-268 on chromosome 1. The yellow rectangle indicates the location of the *Hopscotch* insertion and the green represents the *tb1* ORF. A) Ejutla A; B) Ejutla B; C) La Mesa; D) San Lorenzo. The upper triangle above the black diagonal is colored based on the *r*^2^ value between SNPs while the bottom triangle is colored based on p-value for the corresponding *r*^2^ value.

**Table 3.** mean *r*^2^ values between SNPs on chromosome 1, in the broad *tb1* region, within the 5’ UTR of *tb1* (Region 1), and 66 kb upstream of *tb1* (Region 2).

| Population | Chr. 1 | *tb1* region | Region 1 | Region 2 |
| --- | --- | --- | --- | --- |
| Ejutla A | 0.095 | 0.050 | 0.747 | 0.215 |
| Ejutla B | 0.069 | 0.051 | 0.660 | 0.186 |
| La Mesa | 0.070 | 0.053 | 0.914 | 0.766 |
| San Lorenzo | 0.101 | 0.067 | 0.912 | 0.636 |

The lack of clustering of *Hopscotch* genotypes in our NJ tree as well as the lack of LD around *tb1* do not support the hypothesis that the *Hopscotch* insertion in these populations of *parviglumis* is the result of recent introgression. However, to further explore this hypothesis we performed a STRUCTURE analysis using Illumina MaizeSNP50 data from four of our *parviglumis* populations

(EjuA, EjuB, MSA, and SLO) (Pyhäjärvi et al., 2013) and the maize 282 diversity panel (Cook et al., 2012). The linkage model implemented in STRUCTURE can be used to identify ancestry of blocks of linked variants which would arise as the result of recent admixture between populations. If the *Hopscotch* insertion is present in populations of *parviglumis* as a result of recent admixture with domesticated maize, we would expect the insertion and linked variants in surrounding sites to be assigned to the “maize” cluster in our STRUCTURE runs, not the “teosinte” cluster. In all runs, assignment to maize in the *tb1* region across all four *parviglumis* populations is low (average 0.017) and much below the chromosome-wide average (0.20; Table 4; Fig. 3).

**Figure 3.**
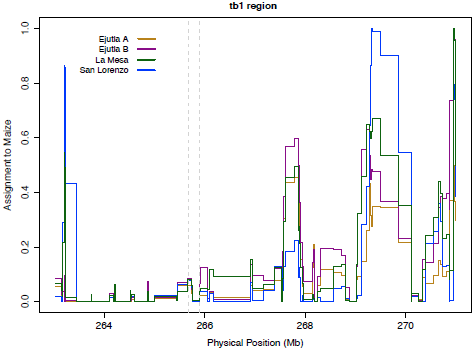
STRUCTURE assignment to maize across a section of chromosome 1. The dotted lines mark the beginning of the sequenced region 66 kb upstream (Region 2) and the end of the *tb1* ORF.

**Table 4.**
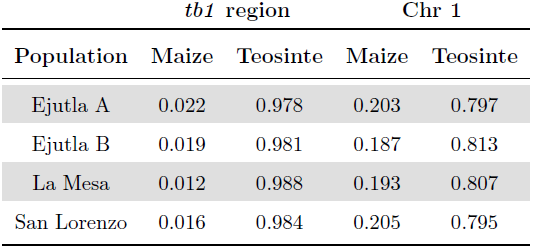
Assignments to maize and teosinte in the *tb1* and chromosome 1 regions from STRUCTURE

### Phenotyping

To assess the contribution of *tb1* to phenotypic variation in tillering in a natural population, we grew plants from seed sampled from the San Lorenzo population of *parviglumis*, which had a high mean frequency (0.44) of the *Hopscotch* insertion based on our initial genotyping. We measured tiller number and tillering index, the ratio of the sum of tiller lengths to plant height, for 206 plants from within the San Lorenzo population, and genotyped plants for the *Hopscotch* insertion. We also measured culm diameter, a phenotype that differs between maize and teosinte (Briggs et al., 2007) but is not thought to be affected by the *Hopscotch* insertion. Phenotypic data are available at http://dx.doi.org/10.6084/m9.figshare.776926. Our plantings produced 82 homozygotes for the *Hopscotch* insertion at *tb1*, 104 heterozygotes, and 20 homozygotes lacking the insertion; these numbers do not deviate from expectations of Hardy-Weinberg equilibrium. After performing a repeated measures ANOVA between our transformed tillering index data and *Hopscotch* genotype we find no significant correlation between genotype at the *Hopscotch* insertion and tillering index (Fig. 4), tiller number, or culm diameter. Only on day 40 did we observe a weak but statistically insignificant (*r*^2^ = 0.02, *p* = 0.0848) correlation between tillering index and the *Hopscotch* genotype, although in the opposite direction of that expected, with homozygotes for the insertion showing a higher tillering index.

**Figure 4.**
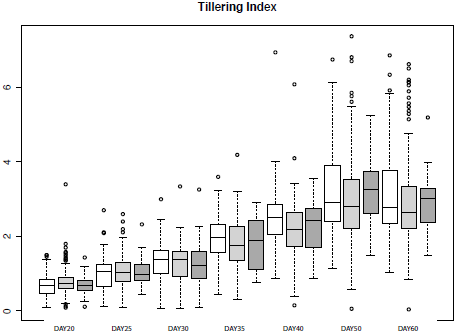
Box-plots showing tillering index in greenhouse grow-outs for phenotyping. White indicates individuals homozygous for the *Hopscotch*, light grey represents heterozygotes, and dark grey represents homozygotes for the teosinte (No *Hopscotch*) allele. Within boxes, dark black lines represent the median, and the edges of the boxes are the first and third quartiles. Outliers are displayed as dots, the maximum value excluding outliers is shown with the top whisker, while the minimum excluding outliers is shown with the bottom whisker.

## DISCUSSION

Adaptation occurs due to selection on standing variation or *de novo* mutations. Adaptation from standing variation has been well-described in a number of systems; for example, selection for lactose tolerance in humans (Plantinga et al., 2012; Tishkoff et al., 2007), variation at the *Eda* locus in three-spined stickleback (Kitano et al., 2008; Colosimo et al., 2005), and pupal diapause in the Apple Maggot fly (Feder et al., 2003). Although the adaptive role of standing variation has been described in many systems, its importance in domestication is not as well studied.

In maize, alleles at domestication loci (*RAMOSA1*, Sigmon and Vollbrecht 2010; *barren stalk1*, Gallavotti et al. 2004; and *grassy tillers1*, Whipple et al. 2011) are thought to have been selected from standing variation, suggesting that diversity already present in teosinte may have played an important role in maize domestication. The *teosinte branched1* gene is one of the best characterized domestication loci, and, while previous studies have suggested that differences in plant architecture between maize and teosinte are a result of selection on standing variation at this locus (Clark et al., 2006; Studer et al., 2011), much remains to be discovered regarding natural variation at this locus and its ecological role in teosinte.

Studer et al. (2011) genotyped 90 accessions of teosinte (inbred and outbred), providing the first evidence that the *Hopscotch* insertion is segregating in teosinte. Given that the *Hopscotch* insertion has been estimated to predate the domestication of maize, it is not surprising that it can be found segregating in populations of teosinte. However, by widely sampling across teosinte populations our study provides greater insight into the distribution and prevalence of the *Hopscotch* in teosinte. While our findings are consistent with Studer et al. (2011) in that we identify the *Hopscotch* allele segregating in teosinte, we find it at higher frequency than previously suggested. The *Hopscotch* allele is more prevalent in *parviglumis* than in *mexicana* in our sample (Appendix 1, See Supplemental Materials with the online version of this article), suggesting a different history of the allele amongst teosinte subspecies. Moreover, many of our *parviglumis* populations with a high frequency of the *Hopscotch* allele fall in the Jalisco cluster identified by Fukunaga et al. (2005), and further distinguish this region from the Balsas River Basin where maize was domesticated (Matsuoka et al., 2002). Potential explanations for the high frequency of the *Hopscotch* element in *parviglumis* from the Jalisco cluster include gene flow from maize, genetic drift, and natural selection.

While gene flow from crops into their wild relatives is well-known, (Ellstrand et al., 1999; Zhang et al., 2009; Thurber et al., 2010; Baack et al., 2008; Hubner et al., 2012; Wilkes, 1977; van Heerwaarden et al., 2011; Barrett, 1983), our results do not suggest introgression from maize at the *tb1* locus, and are more consistent with Hufford et al. (2013) who found resistance to introgression from maize into *mexicana* around domestication loci. Clustering in our NJ trees does not reflect the pattern expected if maize alleles at the *tb1* locus had introgressed into populations of teosinte. Moreover, there is no signature of elevated LD in the *tb1* region relative to the rest of chromosome 1, and Bayesian assignment to a maize cluster in this region is both low and below the chromosome-wide average (Fig. 3, Table 4). Together, these data point to an explanation other than recent introgression for the high observed frequency of *Hopscotch* in the Jalisco cluster of our *parviglumis* populations.

Although recent introgression seems unlikely, we cannot rule out ancient introgression as an explanation for the presence of the *Hopscotch* in these populations. If the *Hopscotch* allele was introgressed in the distant past, recombination may have broken up LD, a process that would be consistent with our data. We find this scenario less plausible, however, as there is no reason why gene flow should have been high in the past but absent in present-day sympatric populations. In fact, early generation maize-teosinte hybrids are common in these populations today (MB Hufford, pers. observation), and genetic data support ongoing gene flow between domesticated maize and both *mexicana* and *parviglumis* in a number of sympatric populations (Hufford et al., 2013; Ellstrand et al., 2007; van Heerwaarden et al., 2011; Warburton et al., 2011).

Remaining explanations for differential frequencies of the *Hopscotch* among teosinte populations include both genetic drift and natural selection. Previous studies using both SSRs and genome-wide SNP data have found evidence for a population bottleneck in the San Lorenzo population (Hufford, 2010; Pyhäjärvi et al., 2013), and the lower levels of sequence diversity in this population in the 5’ UTR (Region 1) coupled with more positive values of Tajima’s D are consistent with these earlier findings. Such population bottlenecks can exaggerate the effects of genetic drift through which the *Hopscotch* allele may have risen to high frequency entirely by chance. A bottleneck in San Lorenzo, however, does not explain the high frequency of the *Hopscotch* in multiple populations in the Jalisco cluster. Moreover, available information on diversity and population structure among Jaliscan populations (Hufford, 2010; Pyhäjärvi et al., 2013) is not suggestive of recent colonization or other demographic events that would predict a high frequency of the allele across populations. Finally, diversity values in the 5’ UTR of *tb1* are suggestive of natural selection acting upon the gene in populations of *parviglumis*. Overall nucleotide diversity is 76% less than seen in the sequences from the 66 kb upstream region, and Tajima’s D is considerably lower and consistently negative across populations (Table 2). In fact, values of Tajima’s D in the 5’ UTR are toward the extreme negative end of the distribution of this statistic previously calculated across loci sequenced in *parviglumis* (Wright et al., 2005; Moeller et al., 2007). Though not definitive, these results are consistent with the action of selection on the upstream region of *tb1*, perhaps suggesting an ecological role for the gene in Jaliscan populations of *parviglumis*.

Significant effects of the *Hopscotch* insertion on lateral branch length, number of cupules, and tillering index in domesticated maize have been well documented (Studer et al., 2011). Weber et al. (2007) described significant phenotypic associations between markers in and around *tb1* and lateral branch length and female ear length in a sample from 74 natural populations of *parviglumis* (Weber et al., 2007); however, these data did not include markers from the *Hopscotch* region 66 kb upstream of *tb1*. Our study is the first to explicitly examine the phenotypic effects of the *Hopscotch* insertion across a wide collection of individuals sampled from natural populations of teosinte. We have found no significant effect of the *Hopscotch* insertion on tillering index or tiller number, a result that is discordant with its clear phenotypic effects in maize. One interpretation of this result would be that the *Hopscotch* controls tillering in maize (Studer et al., 2011), but tillering in teosinte is affected by variation at other loci. Consistent with this interpretation, *tb1* is thought to be part of a complex pathway controlling branching, tillering and other phenotypic traits (Kebrom and Brutnell, 2007; Clark et al., 2006).

A recent study by Studer and Doebley (2012) examined variation across traits in a three-taxa allelic series at the *tb1* locus. Studer and Doebley (2012) introgressed nine unique teosinte *tb1* segments (one from *Zea diploperennis*, and four each from *mexicana* and *parviglumis*) into an inbred maize background and investigated their phenotypic effects. Phenotypes were shown to cluster by taxon, indicating *tb1* may underlie morphological diversification of *Zea*. Additional analysis in Studer and Doebley (2012) suggested tillering index was controlled both by *tb1* and loci elsewhere in the genome. Clues to the identity of these loci may be found in QTL studies that have identified loci controlling branching architecture (*e.g.,* Doebley and Stec 1991, 1993).

Many of these loci (*grassy tillers*, *gt1*; *tassel-replaces-upper-ears1*, *tru1*; *terminal ear1*, *te1)* have been shown to interact with *tb1* (Whipple et al., 2011; Li, 2012), and both *tru1* and *te1* affect the same phenotypic traits as *tb1* (Doebley et al., 1995). *tru1*, for example, has been shown to act either epistatically or downstream of *tb1*, affecting both branching architecture (decreased apical dominance) and tassel phenotypes (shortened tassel and shank length and reduced tassel number; Li 2012). Variation in these additional loci may have affected tillering in our collections and contributed to the lack of correlation we see between *Hopscotch* genotype and tillering.

In conclusion, our findings demonstrate that the *Hopscotch* allele is widespread in populations of *parviglumis* and *mexicana* and occasionally at high allele frequencies. Analysis of linkage using SNPs from across chromosome 1 does not suggest that the *Hopscotch* allele is present in these populations due to recent introgression, and it seems unlikely that the insertion would have drifted to high frequency in multiple populations. We do, however, find preliminary evidence of selection on the *tb1* locus in *parviglumis*. Coupled with our observation of high frequency of the *Hopscotch* insertion in a number of populations, this suggests that the locus may play an ecological role in teosinte.

In contrast to domesticated maize, the *Hopscotch* insertion does not appear to have a large effect on tillering in a diverse sample of *parviglumis* from a natural population and the phenotypic consequences of variation at *tb1* thus remain unclear. Future studies should examine expression levels of *tb1* in teosinte with and without the *Hopscotch* insertion and further characterize the effects of additional loci involved in branching architecture (e.g. *gt1*, *tru1*, and *te1)*. These data, in conjunction with more exhaustive phenotyping, should help to further clarify the ecological significance of the domesticated *tb1* allele in natural populations of teosinte.

## Acknowledgements

The authors thank the Department of Plant Sciences at UC Davis for graduate student research funding to LEV and for research funds supporting the project, UC Mexus for a postdoctoral scholar grant to MBH and JR-I, and G. Coop for helpful discussion.

